# Scalable spike sorting across thousands of neurons by modeling neural dynamics with NeuroSort

**DOI:** 10.1101/2025.11.11.687825

**Authors:** Ling Liu, Zhengwei Hu, Songming Zhang, Yang Liu, Huilin Jia, Hongji Sun, Shengyi Jia, Xiangdong Sun, Jian K. Liu, Xiaojie Duan, Xiaojian Li

**Affiliations:** Shenzhen We-Linking Medical Technology Co., Ltd., Shenzhen, China; School of Computer Science, University of Birmingham, Birmingham, UK; Department of Biomedical Engineering, College of Future Technology, Peking University, Beijing, China; School of Computer Science, Nanjing University, Nanjing, China; School of Computer Engineering and Control Science, Shenzhen University of Advanced Technology, Shenzhen, China; State Key Laboratory of Multi-organ Injury Prevention and Treatment, Key Laboratory of Mental Health of the Ministry of Education, Guangdong-Hong Kong-Macao Greater Bay Area Center for Brain Science and Brain-Inspired Intelligence, Guangdong-Hong Kong Joint Laboratory for Psychiatric Disorders, Guangdong Provincial Key Laboratory of Psychiatric Disorders, Guangdong Basic Research Center of Excellence for Integrated Traditional and Western Medicine for Qingzhi Diseases, Department of Neurobiology, School of Basic Medical Sciences, Southern Medical University, Guangzhou, China

**Keywords:** Spike sorting, feature extraction, clustering, deep learning

## Abstract

Neural circuit research requires resolving the functional coding mechanisms at the single-neuron scale. However, neural signals captured from extracellular recordings are inherently complex, shaped both by electrode properties and the spatially distributed neural population activity across recording sites. These factors introduce strong signal mixing and waveform variation, making it diffcult for existing spike sorting methods to reliably isolate individual neurons from large-scale recodings. To address this, we develop NeuroSort, which fuses spatiotemporal information through a contrastive learning strategy to adaptively model neural dynamics. NeuroSort is a GPU-enabled, highly efficient spike sorting framework, enabling the isolating of up to thousands of neurons. We validated NeuroSort on in vivo extracellular recordings, including Utah array data from rhesus macaques performing motor tasks and Neuropixels 1.0 probe data from a rat during rest. Moreover, in a recording collected by the 1,024-channel Neuroscroll probe, NeuroSort sorted the largest number of well-isolated neurons from different brain areas while achieving a 2-8 times speedup over other methods. Taken together, NeuroSort generalizes across recording conditions, providing an interpretable and scalable paradigm for whole-brain dynamic analysis.

## 1 Introduction

Mechanistic insights into brain function and related diseases through neuroelectrophysiological techniques are essential for advancing precise therapies and neuromodulation technologies [1–4]. Among these, extracellular recording techniques capture continuous electrical signals that reflect the collective activity of multiple neurons. To extract interpretable information, spike sorting is needed to isolate individual neural discharges from these mixed signals.

In real experiments, spike sorting remains challenging because neural recordings are inherently non-stationary across conditions. In addition to the influences of diverse spike waveforms and firing patterns across neurons [5–7] and the spatial relationships between neurons and electrodes [8], variations in electrode properties further introduce probe-specific artifacts into the recordings. Recent advances in low-impedance electrode materials enable long-term, stable recordings with high spatiotemporal resolution [9], while simultaneously introducing substantially greater challenges to the spike sorting pipeline. Specifically, the dramatic increase in electrode counts from hundreds [10, 11] to thousands [12–14] not only leads to explosive growth in data volume but also introduces increased signal complexity due to severe spatiotemporal interactions. Unlike sparse recordings collected by low-throughput arrays, densely packed neural populations cause overlap of electrical fields in highdensity, high-throughput recordings. The heterogeneous spatial distribution of neural activity across these electrodes further complicates the reliable isolation of single-neuron activity. Consequently, the expanding scale and complexity of data driven by next-generation recording technologies render spike sorting the primary technical bottleneck for downstream neural data analysis.

Machine learning (ML) has been widely used in spike sorting in recent years. Specifically, after spike detection, ML algorithms are used for feature extraction and clustering. Almost all tools, including Kilosort v4 [15, 16], MountainSort v5 [17, 18], and SpyKING CIRCUS [19], extract features using native PCA on spike waveforms, yielding low-dimensional but poorly interpretable representations that fail to capture the spatiotemporal variability of neural activity. Graph-based or density-based clustering [20, 21] in sorting methods sharply increases computational demands when scaling to thousands of electrodes, remain time-consuming even for GPU-accelerated methods [15, 16]. Meanwhile, their divide-and-conquer strategies within clustering perform rough pre-clustering with fixed windows, followed by merging or splitting based on domain expertise, further increasing the running burden. These additional interventions, often highly specialized and heuristic, struggle to cope with the signal complexity arising from diverse spatiotemporal interactions. Consequently, current spike sorting methods still require substantial human supervision and are unable to efficiently process large-scale, high-throughput electrophysiological datasets. These challenges highlight the urgent need for adaptive, interpretable and scalable spike sorting paradigms.

In this study, we developed NeuroSort to address all these challenges. NeuroSort adopts an end-to-end deep learning framework that integrates feature extraction and clustering into a unified optimization pipeline (Fig. 1(a)). Given the collected spike waveforms, firing timestamps, and spatial descriptors defined by probe placement and electrode layout, the pipeline first optimizes the model during training and then infers individual neural activities based on the modeled neural dynamics. To clearly capture the spatiotemporal distinctions among neural activities, the spike encoder in NeuroSort learns multi-source fused representations through contrastive learning. Unlike traditional static waveform-based features, these interpretable representations allow detected spikes across all channels to be processed without local partitioning, rapidly yielding well-isolated neurons through the cluster tree. Moreover, based on the spike generation module, domain knowledge about signal variability induced by spatiotemporal interactions can be incorporated at an early stage, which enhances the encoder’s adaptability to diverse spatiotemporal effects, eliminating the need for multistep interventions after clustering. Together, these designs with novel deep learning algorithms enable the pipeline to operate without manual parameter tuning and to generalize across various electrode distributions and brain regions in multiple animals (Fig. 1(b)). NeuroSort consistently outperforms widely used methods in terms of accuracy, speed, and generalization, with particularly high efficiency in thousand-electrode recordings, providing a reliable tool for large-scale electrophysiological data analysis.

**Fig. 1.**
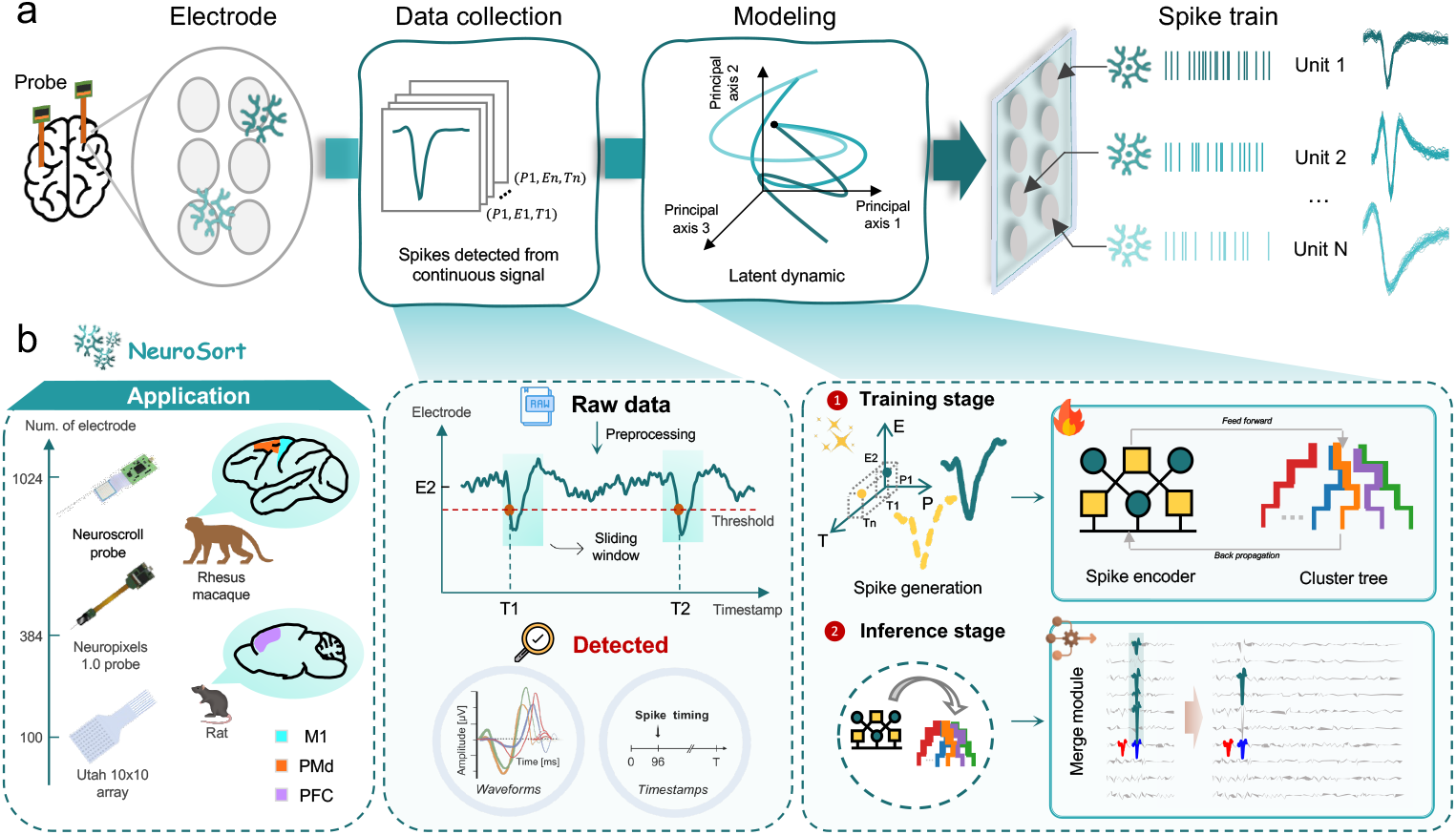
Schematic overview of NeuroSort for spike sorting. (a) Schematic of the pipeline for spike sorting. After preprocessing, spikes are detected using a threshold crossing method. Spikes containing spatiotemporal information are clustered through model training and inference. (b) Application scenarios of NeuroSort, including different electrode types (e.g., Utah array, Neuropixels 1.0 probe, Neuroscroll probe) and various brain structures across multiple animals(e.g., M1 and PMd in rhesus monkeys, PFC in rats, etc.).

## 2 Results

### 2.1 Overview of NeuroSort

NeuroSort is a fully automated spike sorting tool that integrates neural dynamical modeling with a deep learning framework. As a unified deep learning system, NeuroSort comprises three key modules: a spike generator, a shared wide&deep encoder, and a cluster tree (Fig.2). During training, the generator first creates augmented spikes from detected timing, location, and waveform data after preprocesing module. These augmented samples simulate realistic experimental variations, such as those caused by electrode drift, neural bursting, tissue movement, and overlapping signals. By pairing augmented spikes with the original inputs, the wide&deep encoder maps multi-source information into a unified high-dimensional feature space [22, 23] for modeling neural dynamics. The fused representations are then fed into the cluster tree, a tree-structured neural network. It provides a naturally hierarchical architecture for computing assignment probabilities, making it well suited for joint optimization with the encoder under a contrastive learning strategy [24]. During inference, the trained encoder and cluster tree rapidly assign spikes to putative neurons. A final merge module consolidates redundant detections arising from densely packed electrodes.

**Fig. 2.**
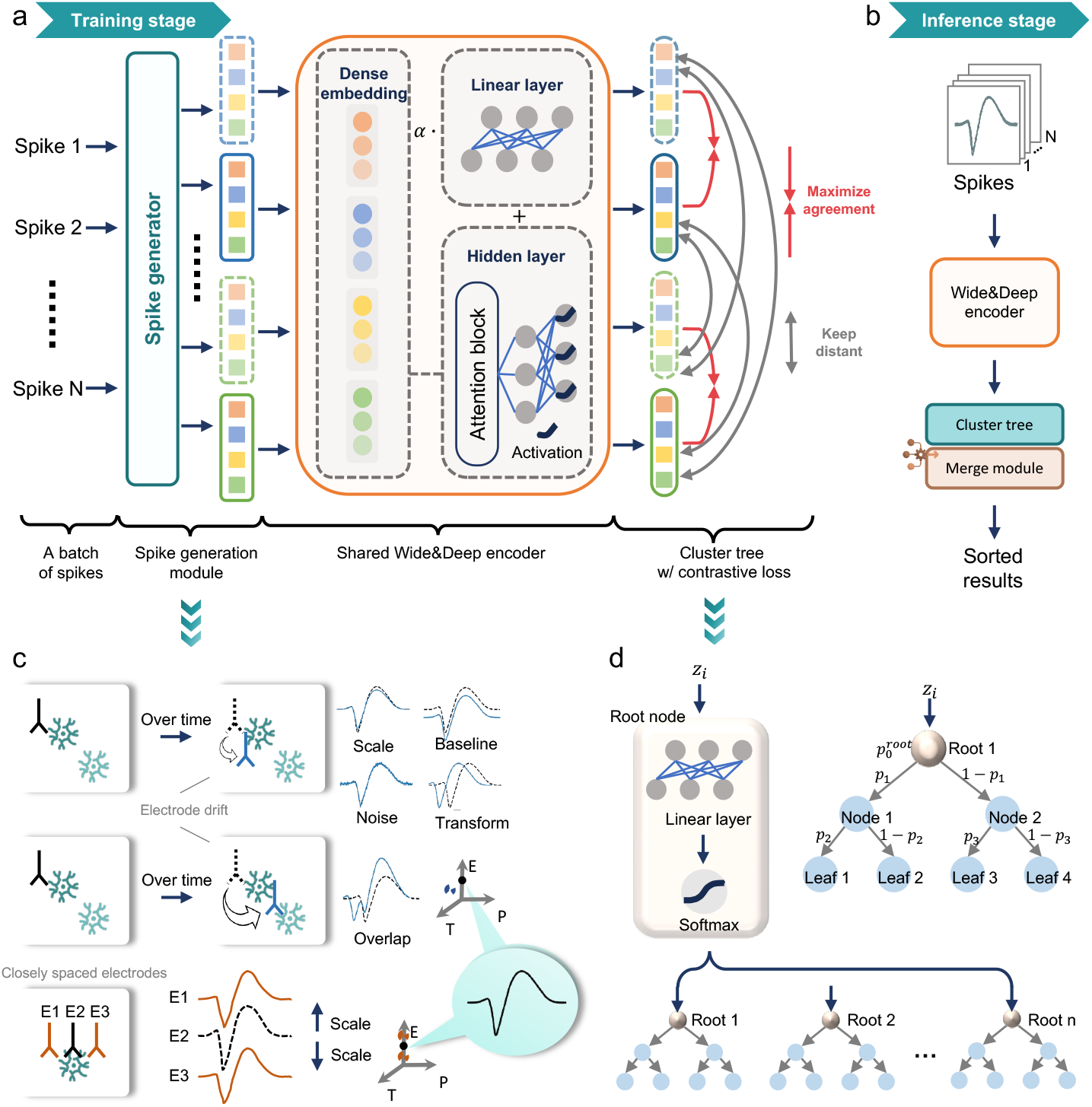
Detailed visualization of NeuroSort framework. (a) Connections between different computational components in NeuroSort during training. (b) Connections between different computational components in NeuroSort during inference. (c) Spike generation module in Neu- roSort. Spatiotemporal augmentations of spikes correspond to potentially detectable activity from the same neuron. The spatiotemporal coordinate system is defined by three dimensions: P (probe), E (electrode), and T (time). In this coordinate system, solid black dots indicate the spatiotemporal information of the input spike, while dashed colored dots indicate the corresponding augmented spikes. In the waveform transformation diagram, the dashed black line and solid colored line represent the waveforms of the input and augmented spikes, respectively. (d) Cluster tree architecture in NeuroSort. The root nodes of the cluster tree are calculated by inputting the latent variables *z*_*i*_ generated by the encoder. The example network shows each tree trained for depth 2.

NeuroSort offers several advantages over previous sorting methods. It provides a fully automated, end-to-end optimized framework and enables fine-grained interpretation of spatiotemporal variability of neural activity. NeuroSort also supports adaptability to diverse recording conditions and facilitates downstream analyses for neuroscientists, including different electrode configurations, brain regions across animals, and experiments spanning multiple spatial and temporal scales. Importantly, NeuroSort delivers high scalability, allowing precise spike sorting in large-scale recordings containing thousands of neurons across multiple regions.

### 2.2 NeuroSort enhances the performance of spike sorting through learning interpretable representations

NeuroSort aims to learn spatiotemporal distinctions among neural activities via its encoder. To assess its effectiveness, we conducted a comparative analysis of the spike features extracted by NeuroSort and those obtained from different spike encoder strategies. For benchmarking, we used extracellular recordings from the primary motor cortex (M1) and dorsal premotor cortex (PMd) of two rhesus macaques [25], each implanted with a 10 × 10 Utah multielectrode array. The M1 and PMd recordings contained 67 and 46 manually labeled single units, respectively.

The features extracted by NeuroSort’s encoder, optimized through contrastive loss, exhibit a clear clustering advantage. We quantified this advantage using the adjusted Rand index (ARI) [26], which measures the consistency between clustering and ground-truth unit labels. As shown in Fig. 3(a), when directly applied to K-means clustering [27], the encoder features achieved an average ARI of 0.68 in M1 recordings. By contrast, removing the contrastive loss, the encoder trained with a reconstruction strategy led to a marked performance decline, with ARI values dropping by about 70.59% (0.68 → 0.20) Replacing the encoder with a linear dimensionality reduction method (PCA) [28] further worsened clustering performance, yielding ARI as low as 0.16. Notably, the ARI remained nearly unchanged (variation ≤ 0.01) even when the PCA feature dimensions were substantially reduced. A similar pattern was observed in recordings from the PMd, confirming that the limitations of reconstruction-based and linear approaches generalize across different cortical areas.

**Fig. 3.**
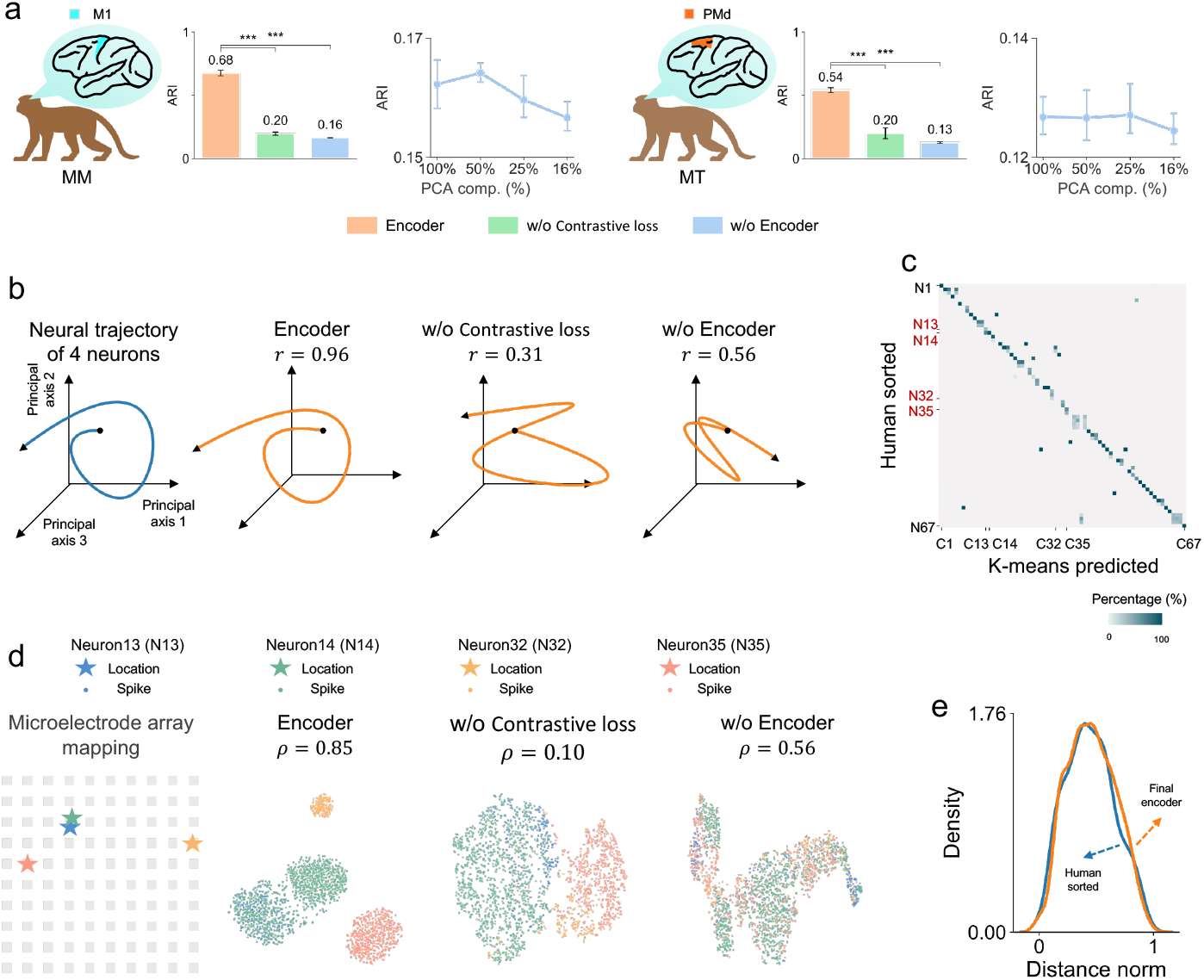
Performance of the NeuroSort’s encoder on extracellular recordings. (a) Quantitative comparison of features extracted with or without the encoder using spikes detected from the motor cortex of two rhesus macaques (MM: M1, left; MT: PMd, right). The ARI was used to evaluate the clustering performance of different feature extraction methods in K-means, as well as that of PCA-based features with varying numbers of principal components. Data are presented as means SD; *p <* 0.001, *p <* 0.01, and *p <* 0.05. (b) Comparison of neural dynamic trajectories derived from the ground-truth population activity (blue) and the latent trajectories learned with or without the encoder (orange). These population dynamics are derived from the four neurons highlighted in red in panel (c). (c) Correspondence between K-means predicted clusters based on encoder features and manually labeled neurons for M1 recordings from rhesus macaque MM, visualized using a pairwise confusion matrix heatmap. The vertical axis represents manually labeled neurons, and the horizontal axis represents predicted clusters. Each matrix entry indicates the proportion of spikes within a predicted cluster that originate from a single manually labeled neuron. (d) Spatial distribution of the four target neurons in the electrode array and their spikes in feature space. Spike features are shown as scatter plots in two-dimensional t-SNE space, with each neuron indicated by a different color. (e) Comparison of the probability density distributions of neuron pair distances based on electrode recording locations (blue) and encoder-based feature distances (orange), estimated using Kernel Density Estimation (KDE).

The superiority of NeuroSort encoder has been further demonstrated in its effective integration of multi-source spike information. We visualized the encoder features from M1 recordings to examine their interpretability on neural dynamics (Fig. 3(b)). Specifically, we randomly selected two spatially neighboring neural subpopulations (spacing *<* 400*µ*m) and two spatially distant subpopulations (spacing *>* 3200*µ*m) in M1 recordings [29, 30]. The encoder features exhibited strong temporal alignment with the neural population dynamics, whereas features derived from reconstruction-based or linear models failed to capture these dynamics. We also evaluated the relationship between the encoder features and the neural population dynamics using the Pearson correlation coefficient [31]. The resulting coefficient reached 0.96, which exceeded the values obtained for the two alternative encoder strategies.

NeuroSort encoder also exhibited clear spatial separation among distinct neurons in the latent space. For this neural population consisting of four neurons (N13, N14, N32, and N35), we visualized the encoder-derived features using t-SNE [32] (Fig. 3(c)), then quantified the correspondence between feature-space distances and physical electrode distances using the Spearman rank correlation coefficient [33] (Fig. 3(d)). The encoder features showed a strong correlation of 0.85 with the true spatial distribution, whereas features derived from other methods did not. Furthermore, across 67 neurons, the distribution of pairwise feature distances from the encoder closely matched the distribution of physical electrode distances, with a Wasserstein distance [34] of 0.016 (Fig. 3(e)). These results indicate that NeuroSort improves spike sorting performance by learning interpretable representations that capture the spatiotemporal structure across neural activities.

### 2.3 NeuroSort demonstrates immense potential to become a superior alternative to human spike sorters

The interpretable representations learned by the NeuroSort encoder provide a foundation for robust tracking of single-neuron activities. Building on this, the cluster tree in NeuroSort further utilizes these representations to adaptively separate individual neurons across varying recording conditions. We continued using M1 and PMd recordings from two rhesus macaques. The performance of the cluster tree was compared against four representative clustering algorithms, including K-means, Gaussian mixture models (GMM), HDBSCAN, and Finch.

In addition to the ARI, we introduced a complementary metric, the Davies–Bouldin Index (DBI) [35], to quantify the compactness and separability of putative neurons (Fig. 4(a)). The cluster tree surpassed other methods in the accuracy of resulting clusters, while also being the most computationally efficient (Fig. 4(b)). Specifically, in M1 recordings, the confusion matrix shows that the majority of sorted spikes fall along the diagonal, indicating that they are consistently assigned according to the manual labels. In contrast, other methods frequently misclassified spikes from nearby neurons, merging them into a single unit while also splitting single neurons into multiple smaller clusters. These inconsistencies, shown as dark off-diagonal blocks in the confusion matrix, underscore the instability of comparative clustering algorithms. When applied to PMd recordings, the cluster tree maintained consistent performance. NeuroSort achieved both high ARI values and low DBI scores, demonstrating the robustness and generalizability of its spike sorting across cortical areas and macaque subjects.

**Fig. 4.**
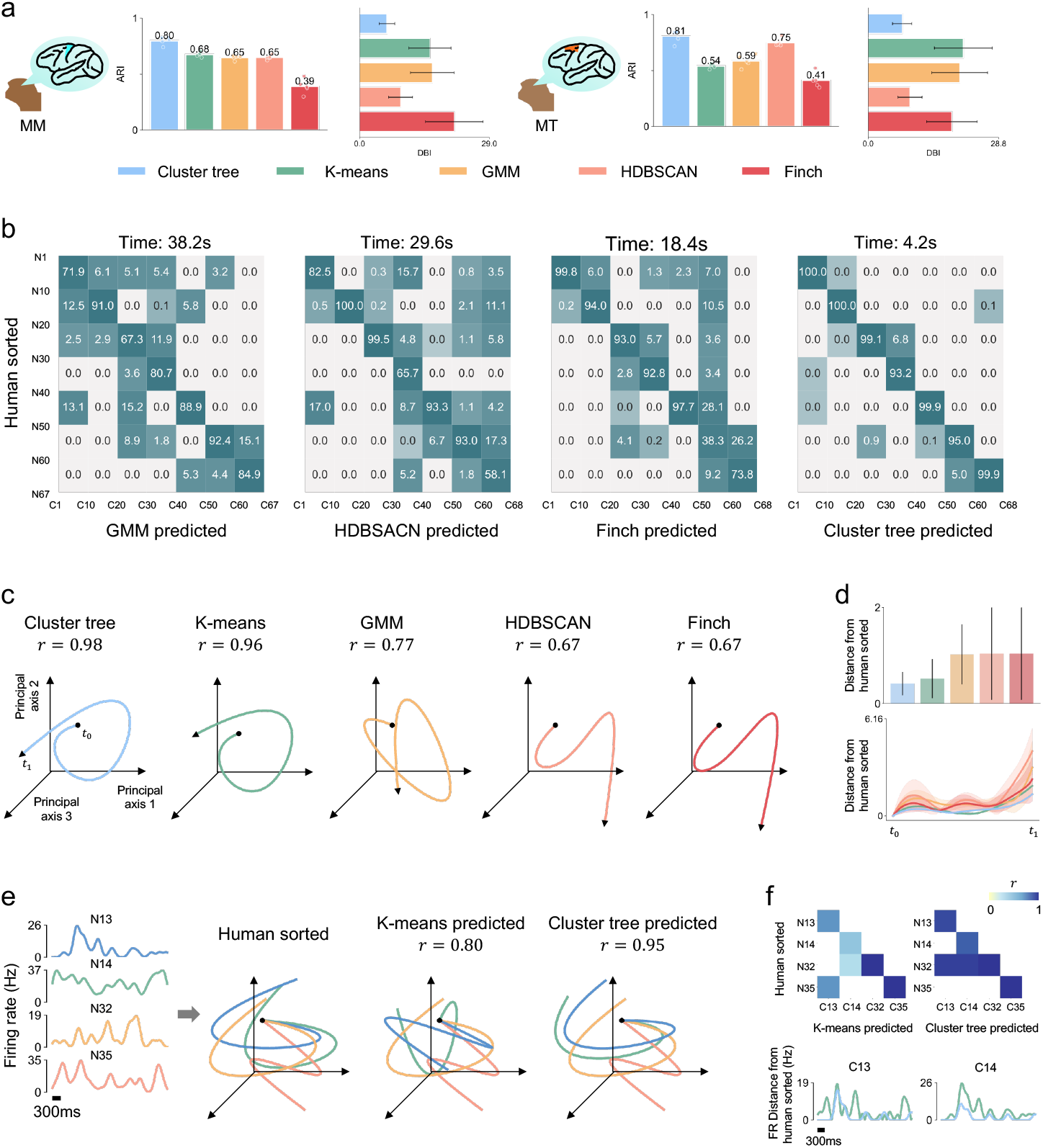
Performance of NeuroSort’s cluster tree on extracellular recordings. (a) Quantitative comparison of clustering performance between the cluster tree and other clustering methods based on encoder features from the motor cortex of two rhesus macaques (MM: M1, left; MT: PMd, right). (b) Comparison of predicted clusters from different clustering methods with manually labeled neurons for M1 recordings of rhesus macaque MM using confusion matrix. The vertical axis represents manually labeled neurons, and the horizontal axis represents predicted clusters. Each matrix entry indicates the proportion of spikes within a predicted cluster originating from a single manually labeled neuron. Average runtime of each method is shown above the corresponding heatmap. (c) Comparison of population dynamic trajectories obtained using different clustering methods. The same color scheme as in panel (a) is used, with blue trajectory showing the population dynamic of the four manually sorted neurons in 3(b). (d) Comparison of population dynamic trajectory distances between inferred and ground-truth neurons, with average values across methods (bar chart) and temporal variations across five independent trials (line chart). (e) Latent trajectories of single neurons within the population are compared between manual and algorithmic inference. (Left) Temporal evolution of firing rates and latent trajectories for manually sorted neurons.(Right) Corresponding trajectories of matched clusters identified by K-means and the cluster tree. (f) Latent trajectory correspondence for four single neurons and firing rate differences for neurons N13 and N14 in panel (e) between manually sorted neurons and clusters inferred by K-means and the cluster tree.

The effectiveness of the cluster tree in NeuroSort can be further demonstrated by its ability to accurately predict neural dynamics (Fig. 4(c)). For the clusters (C13, C14, C32, C35) that most closely matched four manually annotated neurons (N13, N14, N32, N35), the inferred neural trajectories of population dynamics showed the highest correspondence with the ground truth among all compared methods. Specifically, it achieved a Pearson correlation coefficient of 0.98 and the smallest deviation on neural trajectories (Fig. 4(d)). In contrast, K-means clustering exhibited slightly inferior performance, with a 2.04% decrease in correlation and a 25% increase in trajectory deviation. This performance gap arises from the cluster tree’s superior ability to disentangle single-neuron dynamics (Fig.4(e)). Among these four neurons, we observed that the neighboring neurons N13 and N14, along with N32, showed highly similar dynamics (Fig. 4(f)), potentially reflecting their spatial proximity and functional connectivity in the cortex. Consistently, C13 and C14 inferred by the cluster tree displayed strong similarity to N13 and N14, as well as notable similarity to N32 (Fig. 4(g)). However, under K-means clustering, C13 was more similar to N35 than to N32, highlighting K-means’ limitation in capturing the nonlinear relationships encoded among neurons in the cortical circuitry.

### 2.4 NeuroSort significantly outperforms other methods in sorting performance on in vivo recordings

To evaluate NeuroSort’s performance under different recording conditions, we tested a more challenging setup. This dataset employed a Neuropixels 1.0 probe [10] with 384 densely packed electrodes in the prefrontal cortex (PFC) of a Sprague–Dawley (SD) rat. With 384 densely packed electrodes, this configuration created a richly detailed spatiotemporal environment. NeuroSort was benchmarked against other representative automated spike sorting methods, including Kilosort v4 [15] and MountainSort v5 [17], under their default settings. In this PFC recording, the session spanned 32 minutes and captured approximately 10.58 million events.

For each isolated unit, a refractory period of 1 ms was applied, and spike amplitudes were modeled as following a Gaussian distribution to calculate refractory period violations (RPVs) [36] and missed spikes [37]. Typically, units with both RPVs and missed spikes below 5% are considered as “good units”(Fig. 5(a-b)). Compared with Kilosort and MountainSort, NeuroSort identified a higher proportion of good units and contained no units with near-zero signal-to-noise ratio (SNR) [38] (Fig. 5(c)). Across its sorted units, refractory period violations (RPVs) were below 5% for nearly all units, and the number of units with more than 5% missed spikes was the smallest among the compared methods. In contrast, Kilosort and MountainSort exhibit certain limitations. Kilosort produces some low-SNR units, with approximately 3.6% of units having SNR below 4. MountainSort exhibits substantial deficiencies in the completeness of event detection: it isolates far fewer units, approximately 20% of that obtained by Kilosort, and 3.2 times fewer than NeuroSort. Furthermore, among the units it isolates, nearly 40% exhibits a high spike miss rate.

**Fig. 5.**
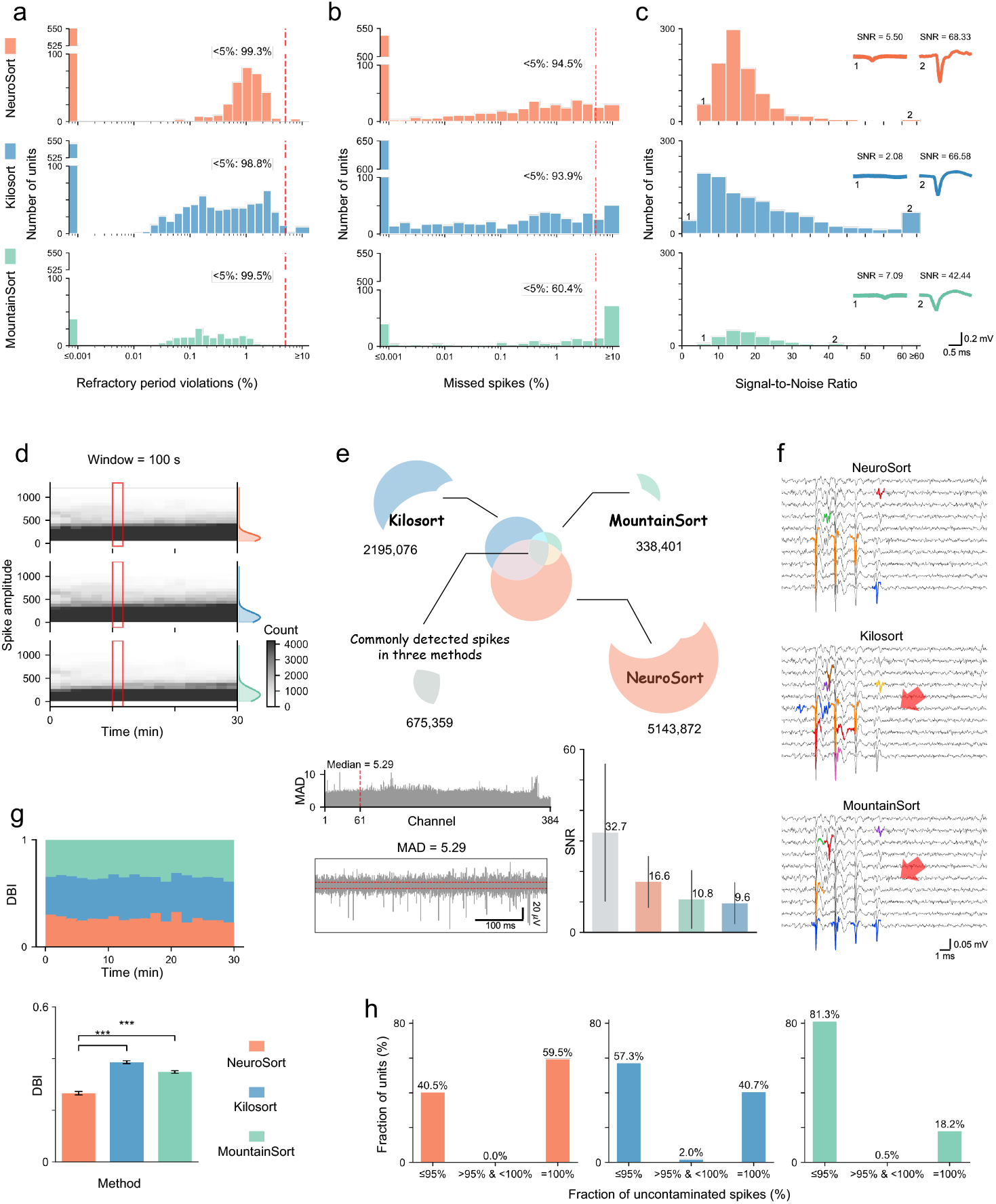
Performance comparison of NeuroSort with other spike sorting methods on 384-channel Neuropixels 1.0 probe recordings from the prefrontal cortex of a Sprague–Dawley (SD) rat. (a) Distribution of the refractory period violations (RPVs) across units in different sorting results. Symbol color indicates the sorting results from NeuroSort (orange), Kilosort (blue), and MountainSort (green). Vertical red dashed lines show the acceptance threshold. (b) Distribution of the proportion of spikes that were missed across units. Vertical red dashed lines show the acceptance threshold. (c) Distribution of signal-to-noise ratio (SNRs) across units. Representative low- and high-SNR units are illustrated, with corresponding template waveforms. (d) Spike amplitude over time during a recording session for different spike sorting methods, along with corresponding amplitude distributions modeled by Gaussian fits. (e) Overlap in spike detection across three methods (top) and corresponding SNRs for concordant and discordant detections (bottom). SNR was computed from the median absolute deviation (MAD) of baseline signals across channels. The median MAD value (5.29) is indicated by the band between the two red dashed lines in the bandpass-filtered signal from channel 61. (f) Representative 20-ms segment illustrating divergent spike sorting results across methods. Red arrows mark waveform regions where other methods diverge from NeuroSort. (g) Comparison of isolated unit quality for commonly detected spikes (gray region in panel (e)). Pairwise t-tests (two-sided, independent) were applied to the Davies–Bouldin Index (DBI) between (i) NeuroSort and KiloSort, and (ii) NeuroSort and MountainSort. Significance levels are denoted as: ∗ ∗ ∗ *p <* 0.001, ∗ ∗ *p <* 0.01, and ∗ *p <* 0.05. (h) Distribution of the fraction of uncontaminated spikes across units in independently detected spikes (orange, blue, and green regions in panel (e)).

The advantages of NeuroSort stem from its broad coverage of spike detection and its high precision in spike identification. In spike detection, NeuroSort can capture more authentic neural events compared with other methods. It identifies substantially more spikes than Kilosort and Mountain- Sort, and these spikes generally exhibit higher amplitudes with lower variance (113. ± 1 87.9 *µ*V for NeuroSort vs. 105.1 ± 113.2 *µ*V for Kilosort) (Fig. 5(d)), suggesting stronger neuronal firing. NeuroSort also recovers a notable proportion of spikes missed by other algorithms. These additional spikes detected show higher signal-to-noise ratios, with median SNR values of 16.6 for NeuroSort compared to 9.6 and 10.8 for Kilosort and MountainSort, respectively (Fig. 5(e)(f)). Moreover, the quality of units separated by NeuroSort is significantly higher than those from Kilosort and MountainSort. For commonly detected spikes, we used the DBI to evaluate the potential misassignment risk of units, and NeuroSort exhibited the lowest risk (Fig. 5(g)). For detected spikes unique to each method, we computed the proportion of uncontaminated spikes in each unit based on RPVs and firing rates (Fig. 5(h)). Under a strict criterion of ≤ 5% contamination, NeuroSort isolated the largest number of units, all free from contamination by other neural activity or noise. This further demonstrates that many spikes missed by other sorting methods were in fact clean neural signals. Overall, NeuroSort demonstrates superior performance in signal stability, isolation quality, and data purity, making it a more reliable tool for neural signal analysis.

### 2.5 NeuroSort effectively processes recordings from up to thousands of electrodes

We further evaluated NeuroSort performance under a larger-scale condition by recording neural activity across multiple cortical and subcortical regions using a Neuroscroll probe with 1,024 channels in the rhesus macaque brain [13]. In this multi-region recording, spanning the premotor areas (F2 and F4), prefrontal area 8A, white matter, striatum (putamen and caudate nucleus), hippocampus, and temporal area 36R, neural activity was recorded for a total of 5 minutes (Fig. 6(a)). We comprehensively evaluated NeuroSort in terms of spike sorting quality, runtime efficiency, and scalability across different hardware configurations relative to other representative methods.

**Fig. 6.**
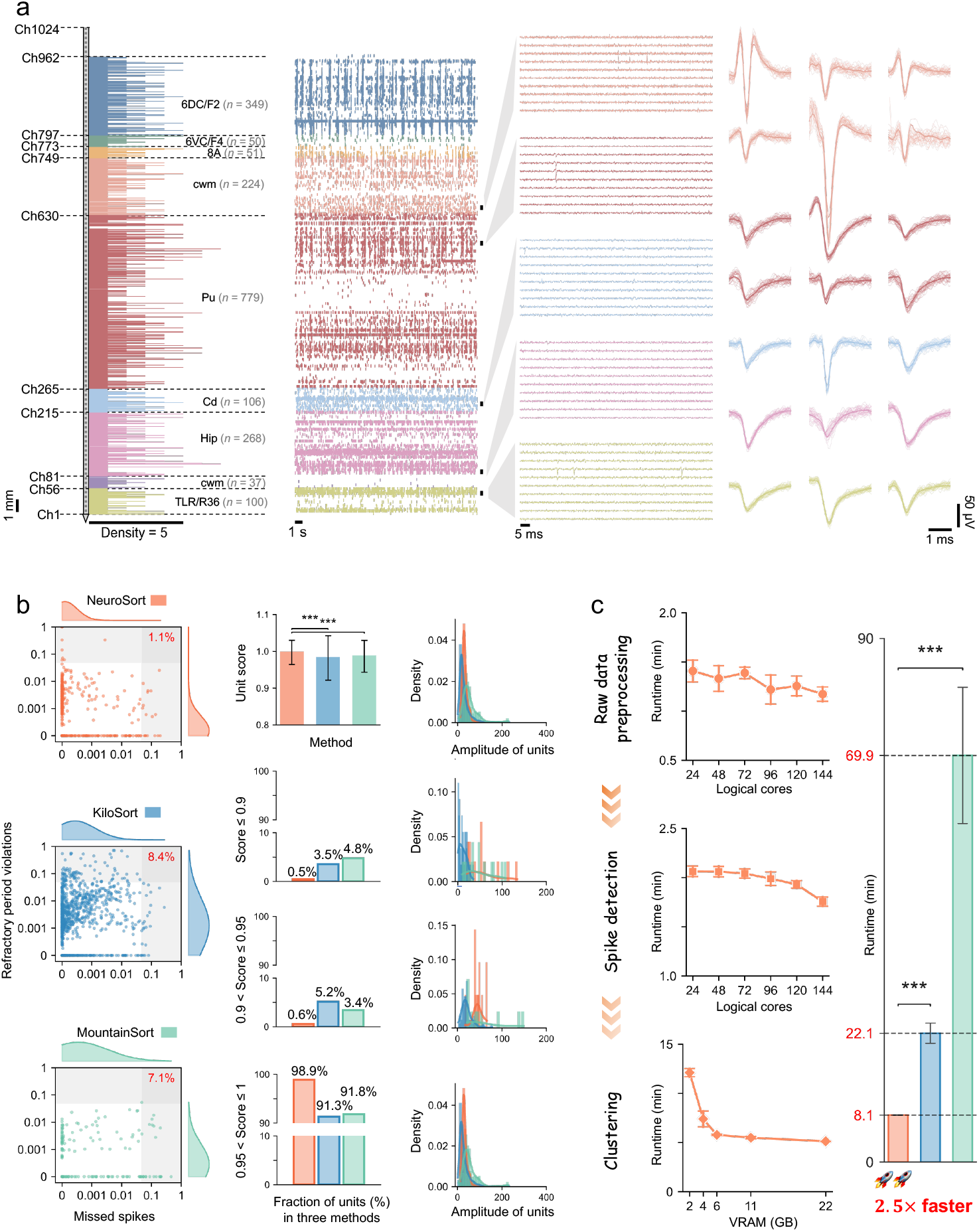
Evaluation of NeuroSort’s performance on cross-region neural recordings from the rhesus macaque acquired with a 1,024-channel Neuroscroll probe. (a) Schematic diagram of NeuroSort’s sorting results. Shown from left to right are the spatial distribution of neurons along the probe track, a 24-second spike train recorded from each channel, a 50-ms AP band trace (spanning five distinct brain structures) from the channels marked by bars, and corresponding representative spike waveforms of different putative units. Spatial distribution of NeuroSort’s sorting results along the probe trajectory in the macaque brain is shown with a schematic of the probe position, the density distribution of putative units across channels, and total units per structure (italicized *n* in parentheses). Symbol color indicates different brain structures. The probe entered the macaque brain from the prefrontal lobe, sequentially passing through area 6 of cortex, dorsocaudal part (Matellis F2) (6DC/F2); area 6 of cortex, ventral part, caudal subdivision (Matellis F4) (6VC/F4); area 8A of cortex (8A); cerebral white matter (cwm); putamen (Pu); caudate nucleus (Cd); hippocampus (Hip); another cwm; area TL, rostral part (area 36R) (TLR/R36). (b) Performance comparison of NeuroSort with Kilosort and MountainSort. Unit isolation quality was evaluated using the sorted score (1 - FP - FN), where FP and FN represent the proportions of RPVs and missed spikes, respectively. Pairwise comparisons of sorted scores were conducted using the Mann–Whitney U test. Significance levels are indicated as: ∗ ∗ ∗ *p <* 0.001, ∗ ∗*p <* 0.01, and ∗ *p <* 0.05. (c) Efficiency comparison of NeuroSort with other spike sorting methods in panel (b). Runtime analysis evaluates the efficiency of NeuroSort’s preprocessing, detection, and clustering modules under different computational resource configurations, as well as its overall runtime compared with Kilosort and MountainSort under default settings. Pairwise runtime comparisons were conducted using two-sided independent t-tests.

NeuroSort detected over 1.3 million events and isolated 1, 964 units. To further strictly assess the quality of each unit, we calculated a sorted score. It is defined as 1 – FP − FN, where FP and FN are the false positive and false negative rates determined by RPVs and missed spikes (Fig. 6(b)). The results show that NeuroSort outperformed all other algorithms, with nearly all units meeting the criteria for good units and an average score of 0.997. Notably, Kilosort produced low-quality units with scores ≤ 0.9 that were concentrated in spikes of smaller amplitude, around 10*µ*V.

In terms of computational efficiency, NeuroSort, substantially outperformed Kilosort and the CPU-only method MountainSort (Fig. 6(c)). Relative to MountainSort, NeuroSort achieved approximately 8× faster runtime when processing over 1.3 million events. It completed the task in 8.1 minutes while operating with a relatively low VRAM usage of 12 GB. And Kilosort required about twice the runtime of NeuroSort and used substantially more memory (22 GB of VRAM). In NeuroSort, computational efficiency improvesas more resources are allocated. For example, preprocessing and spike detection times decrease as more threads are allocated. With 24 threads, preprocessing took approximately 85 s and detection required 124 s. When with 144 logical cores, the preprocessing time was significantly shortened, while the detection time dropped to about 80% of that at 24 threads. Moreover, NeuroSort can still ensure efficient deployment under low resources: even with 2 GB of VRAM, NeuroSort only required 12 minutes.

## 3 Discussion

We have presented a new computational method, NeuroSort, for accurate and interpretable spike sorting via an integrated contrastive learning strategy. Within NeuroSort, the innovative deep feature extraction framework not only accurately captures neural dynamics but also effectively represents the spatial patterns of neurons across electrode arrays. Its unique probe encoding function further enables rapid integration of electrophysiological data collected from multiple brain structures and probe types. NeuroSort is benchmarked against other variants and popular sorting methods, including Kilosort v4 and Mountain v5, and has demonstrated superior performance. This establishes NeuroSort as a powerful and unified solution for integrating multiple neuroscientific research objectives that were previously difficult to achieve. Examples include cell type identification based on firing patterns [36, 39], decoding of neural population dynamics [3, 40–42], tracking neurons across days [43], and automatic identification of disease-specific neuronal firing patterns [44, 45].

NeuroSort also demonstrates strong scalability and usability across the entire spike sorting workflow. Without the need for parameter tuning, NeuroSort substantially reduces the manual curation workload across different probe types (e.g., Utah arrays, Neuropixels probes) and brain regions (including motor and prefrontal cortices), owing to its adaptive algorithmic design. Its end-to-end framework automatically converts raw extracellular recordings into well-isolated single units, providing a fully automated and user-friendly spike sorting solution. Moreover, the GPU-accelerated parallel computing architecture allows NeuroSort to process synchronous activity from thousands of neurons efficiently on large-scale datasets, achieving 2-8 times speedup over other existing methods. With high isolation quality, efficient computational modules, and lower resource consumption compared with other popular sorting methods, NeuroSort can directly scale to analyze neural populations numbering in the tens of thousands. These capabilities are critical for the exploration of simultaneous dynamic interaction and hierarchical integration across multiple brain structures [12, 46].

Although NeuroSort has demonstrated strong generalization across diverse conditions, its performance can still be affected by latent or uncontrolled factors inherent to different experimental preparations. These factors include biological and technical variability, such as experimenter handling, animal state, and recording environment. Therefore, we recommend that users continue to check the results in `Phy` [47]. Given the current limitations of NeuroSort, there are several meaningful directions for future extensions. First, continual learning strategies could be integrated into sorting pipelines to enable rapid generalization to real-time analysis of diverse experimental recordings. Besides, dynamic clustering can be applied to periodically update the reference feature set, thereby compensating for probe drift–induced signal changes. Together, these extensions would help to establish consistent neuronal identity across time in long-term implants or electrophysiological recordings affected by significant brain pulsations [48, 49].

## 4 Methods

### 4.1 Algorithms for NeuroSort

In the following sections, we provide a precise description of the algorithmic procedures employed in NeuroSort.

#### Preprocessing

We designed a stepwise preprocessing pipeline to reduce the impact of artifacts. The pipeline consists of two main steps: Common Average Referencing (CAR), bandpass filtering. The preprocessed signals are prepared for subsequent spike detection and clustering.

##### Common average referencing (CAR)

Raw signals in each batch are processed with CAR to remove noise shared across electrodes. This step enhances inter-channel differences by subtracting the mean across time for each channel and then removing the median across channels at each time point.

##### Bandpass filtering

After CAR, the signals are bandpass filtered to suppress local field potentials (LFPs) and high-frequency noise. A third-order Butterworth filter with a passband from 250 Hz to 7000 Hz is applied, covering the main spectrum of spike signals and minimizing distortion near the edges. Filtering is performed efficiently in the frequency domain using the Fast Fourier Transform (FFT).

#### Event Detection

Spike events are detected from the preprocessed signals using a thresholding method based on the Median Absolute Deviation (MAD).

##### Detection procedure

For each electrode, the median of the signal is calculated to estimate the baseline noise level. A voltage threshold is then set as a multiple *µ* of this median. A time point is identified as a spike event if the signal exceeds this threshold and the time interval from the previous event on the same channel is at least *τ* samples. Default values are *µ* = 3 and *τ* = 10.

##### Spike information extraction

For each detected spike, we record the time point, the corresponding channel and probe, and extract a waveform segment for further analysis. Each waveform segment spans 2ms around the detected time point, with 0.5ms preceding and 1.5ms following the time point. Reducing the threshold can capture low-amplitude spikes but increases the total number of detected events and computational cost during clustering.

#### 4.1.1 Spike generation

NeuroSort generates contrastive spikes to enhance the encoder’s ability to learn discriminative features from realistic neural activity. The goal is to simulate realistic variations in spike signals arising from systematic modifications observed in real recordings, such as electrode drift, temporal shifts, noise, overlapping spikes, and baseline fluctuations.

Each detected spike *x* contains spatiotemporal information: (1) *w*: waveform of the spike, (2) *t*: sampling time of the spike, (3) *e*: electrode where the spike is recorded, and (4) *p*: probe where the spike occurs. A contrastive spike *x*^′^ is generated by combining modified versions of such spatiotemporal information. Specifically, *p*^′^ = *p*, as the distance between probes is much larger than the distance between neurons. Electrode positions are perturbed by a spatial factor ν ∈ {− 1, 0, 1 }: *e*^′^ = *e* + ν. The temporal augmentation is defined as *t*^′^ = *t* + *n*, where *n* is randomly sampled from the interval between the refractory period (0.001 s × sampling rate) and the total recording duration in samples when *e*^′^ = *e*, and set to 0 when *e*^′^ ≠ *e*.

The waveform *w*^′^ is generated using the following approaches:

- **Amplitude scaling**. To account for electrode drift, the waveform is scaled as

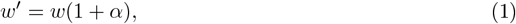

where *α* ∼ *U* (− 0.02, 0.02) for within-channel variations and *α* ∼ *U* (− 0.4, 0.4) for cross-channel variations. This minimizes the impact of amplitude variability from the same neuron on classification.
- **Temporal shifting**. The waveform is shifted along the time axis by *m* ∈ {− 3, − 2, − 1, 1, 2, 3} samples to prevent the constraints of a strict detection window. This can be formally defined from *w* to *w*^′^:

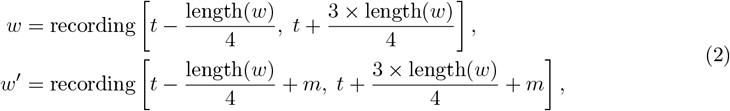

where recording[·] denotes preprocessed recording
- **Noise perturbation**. Multiplicative noise δ ∼ U (0.98, 1.02) is applied:

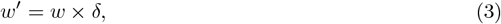

enhancing robustness against noise from microelectrode movement.
- **Overlapping spikes**. For near-simultaneous firing on neighboring electrodes, the waveform *w* is superimposed with a neighboring spike 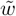:

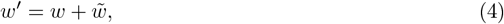

enabling the model to learn features of overlapping signals.
- **Baseline drift**. To simulate baseline fluctuations over long-term recordings, a small additive perturbation *κ* ∼ U (−0.03, 0.03) is applied:

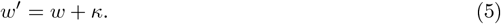

These operations collectively generate contrastive spikes *x*^′^ = (*w*^′^, *t*^′^, *e*^′^, *p*^′^) that reflect realistic variability in neural recordings and provide effective input for the model based on contrastive learning.

#### 4.1.2 Represntation learning via wide & deep encoder

The wide & deep encoder in NeruoSort consists of the following modules:

The wide component is a single linear layer applied directly to the input waveforms *W* to capture fine-grained details of spikes:

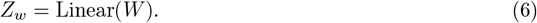

The deep component first embeds discrete features *X* = (*W, T, E, P*) into continuous dense vectors with embedding layers:

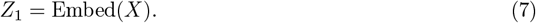

These embeddings, each of dimension 256, are summed and then processed through a 3-layer MLP with multi-head self-attention (MHSA) to model complex spatiotemporal correlations:

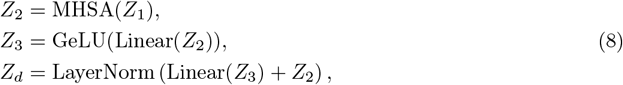

where the MHSA layer employs 8 heads and a dropout rate of 0.1 on attention probabilities. The Linear(·) operation applies position-wise transformations with GeLU activations [50]. Residual connections and layer normalization are applied in the last MLP. The output representations of the wide and deep components are fused with:

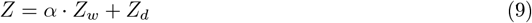

where *α* is a learnable coefficient, initialized to 0.2, that balances contributions from the wide and deep components. The fused feature *Z* has a dimension of 128.

*Z* is fed into a subsequent neural network for clustering, and the encoder is optimized accordingly based on the clustering objective. The encoder and cluster tree constitute a tightly coupled end-to-end system, trained together to encourage the encoder to learn representations that accurately reflect the spatiotemporal variability of neural activity.

#### 4.1.3 Clustering via cluster tree

NeuroSort clusters spikes based on their spatiotemporal features. To accelerate clustering while maintaining accuracy, we employ an improved Fast Feedforward Network [51] called cluster tree. This approach replaces traditional linear-cost feedforward networks with a logarithmic-time complexity structure, suitable for large-scale spike sorting. The cluster tree consists of *Tr* binary trees and a total of *C* leaf nodes:

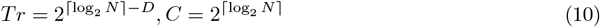

where *N* is the number of electrodes and *D* is the number of tree depth, calculated as:

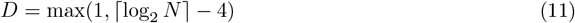

Each node in the cluster tree represents an assignment probability, while leaf nodes act as mixture-of-experts outputs. Feature vectors **z** from the encoder are processed as follows:

- **Root node:** Compute probabilities with a linear transformation followed by softmax: *p*_root_ = softmax

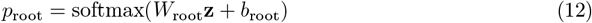
- **Parent nodes:** For each non-root parent node, first apply a linear transformation and ReLU activation:

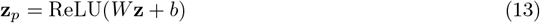

Each *z* in **z**_*p*_ contains two sibling nodes *z*_left_ and *z*_right_. In non-root parent nodes, each pair of sibling nodes, the *z*_left_ and *z*_right_ are used to calculate *p*_left_ and *p*_right_ using StableSoftmax for numerical stability:

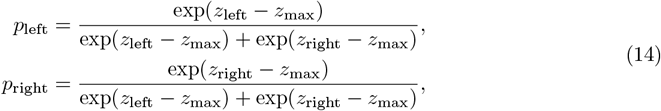

- **Leaf nodes:** Probabilities of leaf nodes are obtained by propagating top-down through the binary tree:

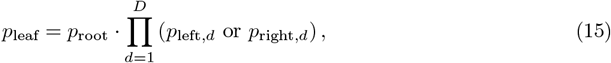

where *p*_left,*d*_ and *p*_right,*d*_ are probabilities of the ancestor node and its sibling at depth *d*. The final prediction of *x* is constructed from all *p*_leaf_ values, forming a probability vector *p*_*x*_.

#### 4.1.4 Joint Training objective

All input spikes can be denoted as *X* = {*x*_1_, *x*_2_, …, *x*_*n*_ }, and their augmented spikes can be represented as *X*^′^ = {*x*^′^ _1_, *x*^′^_2_, …, *x*^′^_*n* }_. The number of leaf nodes *C* corresponds to the number of putative units. The contrastive loss ℒ is formulated in terms of the joint distribution *P* between predicted cluster assignments and its marginals *P*_*c*_ and *P*_*c*_*′* as follows:

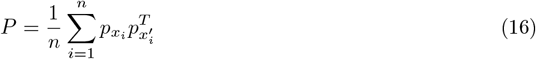

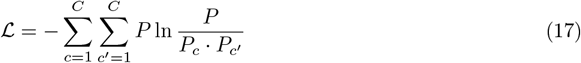

Minimizing ℒ encourages spikes from the same neuron to cluster together, as this maximizes the mutual information between *p*_*x*_ and *p*_*x*_*′* [24]. After convergence, each spike *x*_*i*_ is assigned to the leaf node with the highest probability in 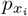. Cluster tree allows efficient, probabilistic clustering of large-scale spikes, combining hierarchical binary tree structure, ReLU-activated parent nodes, StableSoftmax for stability, and mutual information-based training to ensure biologically meaningful clustering.

#### 4.1.5 Eliminating redundant events via merging

After clustering, NeuroSort resolves redundant spikes that arise from temporally overlapping events across multiple channels, which are common in high-density electrode recordings. For each putative unit, spikes occurring within a short temporal window are examined, and only the spike with the largest amplitude is retained as the representative event. In NeuroSort, the merging interval is set to 0.3ms; spikes separated by more than this interval are evaluated in the next merging step. Iterating this procedure across all clusters reconstructs the final, non-redundant spike trains for all putative units.

### 4.2 Performance metrics

#### Clustering metrics

To evaluate clustering quality, we employed two complementary metrics. The Adjusted Rand Index (ARI) [26] provides a supervised assessment by measuring the agreement between the clustering results and the ground-truth units. In contrast, the Davies-Bouldin Index (DBI) [35] provides an unsupervised evaluation of cluster compactness and separation without requiring ground-truth.

##### ARI

ARI was computed by clustering the spike representations produced by the encoder and comparing the resulting clusters with the ground-truth units. The value range of ARI is [− 1, 1], where 1 indicates a perfect match, 0 indicates a random match, and negative values indicate classification performance below chance. Let *N* denotes the total number of spikes, *a*_*i*_ denotes the number of spikes in ground-truth unit *i, b*_*j*_ denotes the number of spikes in putative unit *j*, and *n*_*ij*_ denotes the number of spikes shared between unit *i* and *j*. ARI is defined as:

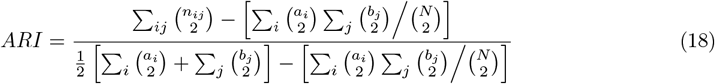

##### DBI

We also used DBI to evaluate the potential misclassification of each putative unit. DBI quantifies isolation quality by comparing the average intra-unit dispersion with the inter-unit separation from its nearest neighboring unit. Let *C* denote the total number of putative units, *C*_*i*_ denotes the number of spikes in unit *i*, and *d*_*ij*_ denotes the Euclidean distance between units *i* and *j*. The intra-unit dispersion *σ*_*i*_ is defined as

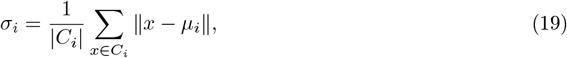

where *µ*_*i*_ is the cluster centroid of unit *i*. DBI is then computed as

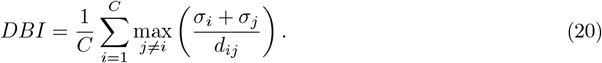

Lower DBI values indicate more compact clusters and better separation between units.

#### Neural dynamics and topology metrics

To interpret the effective representations extracted by the encoder, we assessed them from the perspective of neural population dynamics and topology.

##### Population dynamics similarity

This metric quantifies how closely the temporal patterns of the latent features match the ground-truth neural trajectories. Spike trains of ground-truth units were first summarized over time to form firing rate matrices. Let *X* ∈ ℝ^*T*×*N*^ denote the matrix of groundtruth spike rates, where *T* is the number of time points and *N* is the number of units. Similarly, the latent features extracted by the encoder were summarized into a matrix *Z* ∈ ℝ^*T* × *N*^. Both matrices were then projected into low-dimensional trajectories using Principal Component Analysis (PCA), denoted as *X*_traj_ and *Z*_traj_. The trajectories were aligned using the Kabsch algorithm, and the similarity was quantified as the Pearson correlation coefficient between *X*_traj_ and *Z*_traj_ [29–31].

##### Topological similarity

This metric assesses whether the spatial arrangement of neurons is preserved in the latent feature space. It is computed as the Spearman rank correlation between the distances of cluster centers in the feature space and the distances of the corresponding electrodes [33].

##### Distribution distance

This metric evaluates the overall similarity of pairwise distance distributions between electrodes and the corresponding cluster centers in the feature space. The difference is quantified using the Wasserstein distance [34], reflecting the fidelity of spatial topological encoding.

#### Unit quality metrics

To evaluate the quality of putative units, we computed several popular spike sorting metrics.

**False-positive rate (FP)** were defined as spikes that fell within the refractory period of a unit (± 1 ms from a given spike) and termed refractory period violations (RPVs) [36]. The proportion of false positives was estimated as the quotient between the RPV rate and the mean firing rate [52].

**False-negative rate (FN)** was defined as spikes that were not detected because they fell below the noise threshold of the recording. The proportion of false negatives was estimated by fitting each unit’s spike amplitude distribution with a Gaussian function and quantifying the fraction of area under the curve clipped at the noise threshold [37]. Putative units with a proportion of false positive and false negative rate each below 5% were deemed acceptable.

**Signal-to-noise ratio (SNR)** was defined as the peak absolute voltage of the template across all channels divided by the baseline standard deviation of the same channel, estimated from the median absolute deviation [38]. Units with higher SNRs tend to be more reliable, as the spike amplitude greatly exceeds background fluctuations.

We also used a more stringent metric, termed **sorted score**, to quantify unit isolation quality. The isolation score is defined as 1 − *FP* − *FN*, where *FP* and *FN* represent the false-positive and false-negative rates, respectively.

To quantify the degree of contamination in the spikes detected by each method, we defined a **contamination score** as

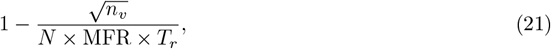

where *n*_*v*_ is the number of refractory period violations, *N* is the total number of spikes, MFR is the mean firing rate, and *T*_*r*_ is the refractory period interval, set to 0.001 s. A higher contamination score indicates fewer contaminated spikes and better unit isolation.

#### Efficiency and Scalability

We evaluated NeuroSort’s processing efficiency under different computational resource configurations. For fair comparison, all spike sorting methods were tested under the same settings. The configurations varied in the number of CPU logical cores of 24, 48, 72, 96, 120 and 144, and GPU memory capacities of 2 GB, 4 GB, 6 GB, 11 GB and 22 GB. This setup allowed us to assess both runtime performance and scalability.

### Benchmarking encoder algorithms

To comprehensively assess the effectiveness of the encoder, we compared three encoder strategies: (1) the full encoder, (2) the encoder without the contrastive learning strategy, and (3) directly using PCA features (without encoder). All learnt features were used as input for K-Means clustering to evaluate unit quality.

The **full encoder** corresponds to the encoder module designed in NeuroSort, which incorporates contrastive learning to enhance the discriminability of spike representations. Details of its structure and training objective are described in Section 4.1.

The **encoder without the contrastive learning strategy** employs a Contractive Autoencoder (CAE) [53]. The CAE learns a mapping from the input spike waveform to a latent representation and reconstructs the original waveform, while applying a Jacobian regularization term. The loss function of CAE is defined as:

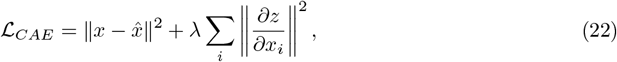

where the first term denotes the reconstruction loss and the second term enforces the contractive constraint. The hyperparameter *λ* was typically set to 0.5

The **directly using PCA features strategy** directly applies PCA to the spike waveforms, reducing their dimensionality without involving any deep learning modules.

### 4.4 Benchmarking clustering algorithms

To test the performance of the proposed cluster tree module in NeuroSort, we compared it with four clustering algorithms: K-Means, Gaussian Mixture Model (GMM) [54], HDBSCAN [55], and Finch [56]. Each method was evaluated on the same set of spike features learnt by the encoder.

#### K-Means

After specifying the cluster number, K-Means [27] iteratively assigns each spike to its nearest cluster center and updates the centroids until convergence. The algorithm minimizes the overall intra-cluster distance between spikes and their assigned centers, returning the cluster index for each spike.

#### Gaussian Mixture Model (GMM)

Similar to K-Means, GMM requires specifying the number of clusters in advance. However, instead of hard assignments, it performs soft clustering by modeling the data as a mixture of Gaussian distributions. The parameters of these distributions are estimated through the Expectation-Maximization (EM) algorithm, which alternates between estimating cluster probabilities for each spike and updating the Gaussian parameters until convergence.

#### HDBSCAN

HDBSCAN performs hierarchical clustering based on a Minimum Spanning Tree (MST), allowing detection of neurons with varying spike densities. The key parameters `min_sa mples` and `min_cluster_size` control the minimum number of neighboring spikes defining a dense region and the minimum cluster size, respectively. To align the cluster count with the number of ground truth neurons, we set `min_samples` to 1 and `min_cluster_size` to 10. To accelerate clustering, the features output from the encoder are first reduced to a compact two-dimensional space using UMAP before HDBSCAN clustering [57].

#### Finch

Finch performs hierarchical clustering without requiring the number of clusters as input. It constructs a nearest neighbor graph and progressively merges samples based on connectivity, discovering spike clusters at multiple levels of granularity. Each layer represents a different clustering resolution, and we selected the layer whose cluster number most closely matched the number of ground truth neurons for evaluation.

### 4.5 Benchmarking spike sorting algorithms

To evaluate the performance of automated spike sorting methods, we tested NeuroSort together with two representative algorithms, Kilosort v4 [15] and MountainSort v5 [17]. The evaluation was performed on electrophysiological data collected from the Neuropixels 1.0 probe [10] and the Neuroscroll probe [13]. All algorithms were executed in fully automated mode using their respective default parameter settings.

#### Kilosort v4

Kilosort is a widely adopted spike sorting family based on template matching and drift correction, originally introduced for high-channel count probes. The first version implemented a GPU-accelerated matching pursuit algorithm, while subsequent versions (Kilosort v2 and Kilosort v3) progressively improved robustness to sorting performance. Kilosort v4 extends this framework by incorporating a graph-based clustering approach, enabling more accurate spike detection in long-duration recordings with substantial tissue movement.

#### MountainSort v5

MountainSort is an open-source spike sorter designed for transparency and reproducibility. Earlier versions (MountainSort v1–4) used density-based clustering and local principal component features, while MountainSort v5 introduces a modular and scalable architecture optimized for modern highdensity probes. The new version integrates automated noise suppression, template merging, and drift compensation modules, achieving improved sorting accuracy with minimal parameter tuning.

### 4.6 Datasets description

#### Utah array dataset

We used the publicly available extracellular dataset with known ground truth collected by Matthew G. Perich (https://crcns.org/data-sets/motor-cortex/pmd-1/about-pmd-1). The recordings were obtained with 10 × 10 Utah multielectrode arrays (Blackrock Microsystems, Salt Lake City, UT) arranged on a square grid with 400*µ*m inter-electrode spacing. The dataset includes simultaneous neural activity and behavioral measurements together with ground-truth spike information.

#### Neuropixels 1.0 probe dataset

The Neuropixels 1.0 dataset was collected by Z.H. in our laboratory. All procedures were approved and conducted in accordance with ethical and animal welfare guidelines. In this experiment, a Sprague–Dawley (SD) rat was anesthetized with isoflurane, and a Neuropixels 1.0 probe was implanted in the prefrontal cortex. Approximately 25 minutes after implantation, neural activity was continuously recorded for 32 minutes across 384 recording sites. The probe contains 384 sites arranged in two staggered columns, with a vertical spacing of 20*µ*m, a horizontal spacing 12*µ*m larger than the vertical distance, and a 16*µ*m horizontal offset between adjacent rows.

#### Neuroscroll probe dataset

The dataset was obtained by Liu *et al*. [13] using the Neuroscroll probe interfaced with a 1,024- channel Intan amplifier system. The probe measures 47mm in length, with recording sites evenly spaced across the two columns, and adjacent rows horizontally offset. Electrodes within each column are spaced 72*µ*m apart, while the spacing between columns 44*µ*m. All electrophysiological recordings analyzed in this study were sampled at 30 kHz.

### 4.7 Data formats

#### Input data

NeuroSort takes a binary data file as input. By default, the binary file should use the ‘int16’ data type. If the data type differs, users must explicitly specify one of the following alternatives: ‘uint16’, ‘int32’, or ‘float32’. For data in other formats (e.g., ‘nwb’, ‘open-ephys’, ‘blackrock’, ‘neuralynx’, or ‘intan’), the file must be converted to a compatible binary format before input. `Kilosort` provides example notebooks demonstrating such conversions (https://github.com/MouseLand/Kilosort/blob/main/docs/tutorials/load_data.ipynb).

#### Auxiliary input information

In addition to the binary raw data, NeuroSort needs to use auxiliary information, including channel count, sampling rate, and ADC-to-*µ*V scaling factor. And the electrode map is optional. If channel numbers follow inter-electrode distance, an electrode map is not required, but the channel mode must be set to indicate whether single-neuron activity spans multiple electrodes. Otherwise, an electrode map is required, except when using mutually independent electrodes (typically spaced more than 50 *µ*m apart). In that case, simply set the channel mode directly.

Users can also set custom voltage thresholds for spike detection through the interface, similar to real-time detection provided by `Intan RHX`, enabling direct user–tool interaction. By default, spike detection is fully automated, with spikes on each channel identified through adaptive, programmatic computations. In addition, users can optionally save the preprocessing file and detected spike waveforms. These intermediate files are not saved by default to minimize I/O overhead.

#### Output data

By default, NeuroSort outputs a lightweight ‘.h5’ file containing spike timestamps, corresponding channels, and cluster IDs. In this file, negative cluster IDs indicate temporally overlapping spikes influenced by multiple channels. Although these values are not directly related to neuronal firing analysis, they provide intuitive information for manual supervision of spike sorting quality. Unlike Kilosort v4 and MountainSort v5, which require additional scripts to obtain channel information, NeuroSort integrates this functionality as part of its standard output. The ‘.h5’ file can be further converted into a ‘.npy’ file for visualization in `Phy` [47] using the provided example notebook (https://github.com/NeuroAILand/NeuroSort/main/tutorials/load_result.ipynb). If the user has additionally enabled the option to save preprocessing files or spike waveforms, these intermediate files will also be generated.

### 4.8 The implementation details of NeuroSort

#### Implementation settings

NeuroSort is implemented in Python 3.9. Core computations rely on libraries including `PyTorch-CUDA, NumPy, SciPy, Signal, Threading, Concurrent`, and `h5py`. All data analysis and visualization were also performed in Python 3.9, using `matplotlib` and `seaborn` to generate figures. Filtering during preprocessing uses `signal`, while spike detection uses `SciPy`‘s `find_peaks` following Butterworth band-pass filtering with `filtfilt`. The preprocessing and detection stages run primarily on the CPU, while the clustering module is executed on GPU using `PyTorch`. Memory management is optimized through `gc` for improved access and resource release.

#### Parallel acceleration

NeuroSort is designed with modularity and high performance as its core objectives. The entire pipeline is optimized across preprocessing, detection, and clustering stages to efficiently utilize both system memory (RAM) and GPU memory (VRAM) through parallelism.

For high-throughput datasets, the preprocessing stage adopts a chunk-wise strategy based on data size, while the detection stage operates channel-wise to eliminate boundary effects. Both stages are parallelized using multithreading to maximize CPU utilization. Parallel execution is implemented with `ThreadPoolExecutor` and `as_completed` from the `concurrent` package, and file I/O safety during concurrent operations is ensured with `Lock` from the `threading` module.

In the clustering stage, batch-wise operations are used to further accelerate computation, and the `num_workers` parameter in `PyTorch` is tuned to achieve balanced memory access and thread-level efficiency.

#### Computation setting

All spike sorting methods were run on the same workstation (144-thread Intel Xeon Platinum 8360Y, 503 GiB RAM, NVIDIA GeForce RTX 4090 GPU, and Ubuntu 20.04 LTS).

## 5 Data avaibility

The publicly available monkey data recorded by Utah multielectrode arrays were made available in https://crcns.org/data-sets/motor-cortex/pmd-1/about-pmd-1. The rhesus macaque electrophysiology data used in this study were collected using the Neuroscroll probe in a prior study. The data analyzed during the current study are available at https://github.com/NeuroAILand/NeuroSort. Any additional requests for information, including access to raw data collected in this laboratory, can be directed to the corresponding author upon reasonable request. Source data are provided in this paper.

## 6 Code avaibility

Code for the method developed in this study is available at https://github.com/NeuroAILand/NeuroSort with license details described. For performance comparison, KiloSort 4.0 and MountainSort 5.0 are available at https://github.com/MouseLand/Kilosort and https://github.com/flatironinstitute/mountainsort5, respectively.

## Acknowledgements

We thank Lang Qian, Shengjie Zheng, and Liang Ma for helpful discussions. This research was supported by the National Key Research and Development Program of China (Grant No. 2023YFE0101400) and the Joint Laboratory of Brain–Machine Intelligence Technology in collaboration with Shenzhen We-Linking Medical Technology Co., Ltd.

## Notes

https://github.com/NeuroAILand/NeuroSort

### Competing Interest Statement

The authors have declared no competing interest.

## References

[1] Zeng, Z., Zhang, C., Xu, Y., He, H., Gu, Y.: Distinct neural population code and causal roles of primate caudate nucleus in multimodal decision-making. Nature Communications 16(1), 5253 (2025)

[2] Dimakou, A., Pezzulo, G., Zangrossi, A., Corbetta, M.: The predictive nature of spontaneous brain activity across scales and species. Neuron 113(9), 1310–1332 (2025)

[3] Urai, A.E., Doiron, B., Leifer, A.M., Churchland, A.K.: Large-scale neural recordings call for new insights to link brain and behavior. Nature neuroscience 25(1), 11–19 (2022)

[4] Paulk, A.C., Kfir, Y., Khanna, A.R., Mustroph, M.L., Trautmann, E.M., Soper, D.J., Stavisky, S.D., Welkenhuysen, M., Dutta, B., Shenoy, K.V., et al.: Large-scale neural recordings with single neuron resolution using neuropixels probes in human cortex. Nature neuroscience 25(2), 252–263 (2022)

[5] Van Vreeswijk, C., Sompolinsky, H.: Chaos in neuronal networks with balanced excitatory and inhibitory activity. Science 274(5293), 1724–1726 (1996)

[6] Geva, N., Deitch, D., Rubin, A., Ziv, Y.: Time and experience differentially affect distinct aspects of hippocampal representational drift. Neuron 111(15), 2357–2366 (2023)

[7] Vishne, G., Gerber, E.M., Knight, R.T., Deouell, L.Y.: Distinct ventral stream and prefrontal cortex representational dynamics during sustained conscious visual perception. Cell reports 42(7) (2023)

[8] Gold, C., Henze, D.A., Koch, C., Buzsaki, G.: On the origin of the extracellular action potential waveform: a modeling study. Journal of neurophysiology 95(5), 3113–3128 (2006)

[9] Sheng, H., Liu, R., Li, Q., Lin, Z., He, Y., Blum, T.S., Zhao, H., Tang, X., Wang, W., Jin, L., et al.: Brain implantation of soft bioelectronics via embryonic development. Nature, 1–11 (2025)

[10] Jun, J.J., Steinmetz, N.A., Siegle, J.H., Denman, D.J., Bauza, M., Barbarits, B., Lee, A.K., Anastassiou, C.A., Andrei, A., Aydın, Ç., et al.: Fully integrated silicon probes for high-density recording of neural activity. Nature 551(7679), 232–236 (2017)

[11] Shandhi, M.M.H., Negi, S.: Fabrication of out-of-plane high channel density microelectrode neural array with 3d recording and stimulation capabilities. Journal of Microelectromechanical Systems 29(4), 522–531 (2020)

[12] Trautmann, E.M., Hesse, J.K., Stine, G.M., Xia, R., Zhu, S., O’Shea, D.J., Karsh, B., Colonell, J., Lanfranchi, F.F., Vyas, S., et al.: Large-scale high-density brain-wide neural recording in nonhuman primates. Nature Neuroscience, 1–14 (2025)

[13] Liu, Y., Jia, H., Sun, H., Jia, S., Yang, Z., Li, A., Jiang, A., Naya, Y., Yang, C., Xue, S., et al.: A high-density 1,024-channel probe for brain-wide recordings in non-human primates. Nature Neuroscience 27(8), 1620–1631 (2024)

[14] Steinmetz, N.A., Aydin, C., Lebedeva, A., Okun, M., Pachitariu, M., Bauza, M., Beau, M., Bhagat, J., Böhm, C., Broux, M., et al.: Neuropixels 2.0: A miniaturized high-density probe for stable, long-term brain recordings. Science 372(6539), 4588 (2021)

[15] Pachitariu, M., Sridhar, S., Pennington, J., Stringer, C.: Spike sorting with kilosort4. Nature Methods, 1–8 (2024)

[16] Pachitariu, M., Steinmetz, N.A., Kadir, S.N., Carandini, M., Harris, K.D.: Fast and accurate spike sorting of high-channel count probes with kilosort. Advances in neural information processing systems 29 (2016)

[17] Chung, J.E., Magland, J.F., Barnett, A.H., Tolosa, V.M., Tooker, A.C., Lee, K.Y., Shah, K.G., Felix, S.H., Frank, L.M., Greengard, L.F.: MountainSort5. https://github.com/flatironinstitute/mountainsort5 (2024)

[18] Chung, J.E., Magland, J.F., Barnett, A.H., Tolosa, V.M., Tooker, A.C., Lee, K.Y., Shah, K.G., Felix, S.H., Frank, L.M., Greengard, L.F.: A fully automated approach to spike sorting. Neuron 95(6), 1381–1394 (2017)

[19] Yger, P., Spampinato, G.L., Esposito, E., Lefebvre, B., Deny, S., Gardella, C., Stimberg, M., Jetter, F., Zeck, G., Picaud, S., et al.: A spike sorting toolbox for up to thousands of electrodes validated with ground truth recordings in vitro and in vivo. Elife 7, 34518 (2018)

[20] Magland, J.F., Barnett, A.H.: Unimodal clustering using isotonic regression: Iso-split. arXiv preprint 1508.04841 (2015)

[21] Rodriguez, A., Laio, A.: Clustering by fast search and find of density peaks. science 344(6191), 1492–1496 (2014)

[22] Cheng, H.-T., Koc, L., Harmsen, J., Shaked, T., Chandra, T., Aradhye, H., Anderson, G., Corrado, G., Chai, W., Ispir, M., et al.: Wide & deep learning for recommender systems. In: Proceedings of the 1st Workshop on Deep Learning for Recommender Systems, pp. 7–10 (2016)

[23] Vaswani, A., Shazeer, N., Parmar, N., Uszkoreit, J., Jones, L., Gomez, A.N., Kaiser, Ł., Polosukhin, I.: Attention is all you need. Advances in neural information processing systems 30 (2017)

[24] Ji, X., Henriques, J.F., Vedaldi, A.: Invariant information clustering for unsupervised image classification and segmentation. In: Proceedings of the IEEE International Conference on Computer Vision, pp. 9865–9874 (2019)

[25] Perich, M.G., Lawlor, P.N., Kording, K.P., Miller, L.E., et al.: Extracellular neural recordings from macaque primary and dorsal premotor motor cortex during a sequential reaching task. CRCNS. org 10, 0–872 (2018)

[26] Hubert, L., Arabie, P.: Comparing partitions. Journal of classification 2(1), 193–218 (1985)

[27] Hartigan, J.A., Wong, M.A.: Algorithm as 136: A k-means clustering algorithm. Journal of the royal statistical society. series c (applied statistics) 28(1), 100–108 (1979)

[28] Abdi, H., Williams, L.J.: Principal component analysis. Wiley interdisciplinary reviews: computational statistics 2(4), 433–459 (2010)

[29] Vyas, S., Golub, M.D., Sussillo, D., Shenoy, K.V.: Computation through neural population dynamics. Annual review of neuroscience 43(1), 249–275 (2020)

[30] Trautmann, E.M., Stavisky, S.D., Lahiri, S., Ames, K.C., Kaufman, M.T., O’Shea, D.J., Vyas, S., Sun, X., Ryu, S.I., Ganguli, S., et al.: Accurate estimation of neural population dynamics without spike sorting. Neuron 103(2), 292–308 (2019)

[31] Schober, P., Boer, C., Schwarte, L.A.: Correlation coefficients: appropriate use and interpretation. Anesthesia & analgesia 126(5), 1763–1768 (2018)

[32] Maaten, L., Hinton, G.: Visualizing data using t-sne. Journal of machine learning research 9(Nov), 2579–2605 (2008)

[33] Spearman, C.: The proof and measurement of association between two things. International journal of epidemiology 39(5), 1137–1150 (2010)

[34] Panaretos, V.M., Zemel, Y.: Statistical aspects of wasserstein distances. Annual review of statistics and its application 6(1), 405–431 (2019)

[35] Davies, D.L., Bouldin, D.W.: A cluster separation measure. IEEE transactions on pattern analysis and machine intelligence (2), 224–227 (2009)

[36] Beau, M., Herzfeld, D.J., Naveros, F., Hemelt, M.E., D’Agostino, F., Oostland, M., Sánchez-López, A., Chung, Y.Y., Maibach, M., Kyranakis, S., et al.: A deep learning strategy to identify cell types across species from high-density extracellular recordings. Cell 188(8), 2218–2234 (2025)

[37] Fabre, J.M., Beest, E., Peters, A., Carandini, M., Harris, K.: Bombcell: automated curation and cell classification of spike-sorted electrophysiology data. Zenodo (2023)

[38] Wyngaard, A.J., Llobet, V., Barbour, B.: Lussac: a fully-automated consensus method that increases the yield and quality of spike-sorting analyses. bioRxiv, 2022–02 (2022)

[39] Lui, J.H., Nguyen, N.D., Grutzner, S.M., Darmanis, S., Peixoto, D., Wagner, M.J., Allen, W.E., Kebschull, J.M., Richman, E.B., Ren, J., et al.: Differential encoding in prefrontal cortex projection neuron classes across cognitive tasks. Cell 184(2), 489–506 (2021)

[40] Gosztolai, A., Peach, R.L., Arnaudon, A., Barahona, M., Vandergheynst, P.: Marble: interpretable representations of neural population dynamics using geometric deep learning. Nature Methods, 1–9 (2025)

[41] Karpowicz, B.M., Ali, Y.H., Wimalasena, L.N., Sedler, A.R., Keshtkaran, M.R., Bodkin, K., Ma, X., Rubin, D.B., Williams, Z.M., Cash, S.S., et al.: Stabilizing brain-computer interfaces through alignment of latent dynamics. Nature Communications 16(1), 4662 (2025)

[42] Kim, J.-H., Daie, K., Li, N.: A combinatorial neural code for long-term motor memory. Nature 637(8046), 663–672 (2025)

[43] Beest, E.H., Bimbard, C., Fabre, J.M., Dodgson, S.W., Takács, F., Coen, P., Lebedeva, A., Harris, K.D., Carandini, M.: Tracking neurons across days with high-density probes. Nature Methods 22(4), 778–787 (2025)

[44] Keller, C.J., Truccolo, W., Gale, J.T., Eskandar, E., Thesen, T., Carlson, C., Devinsky, O., Kuzniecky, R., Doyle, W.K., Madsen, J.R., et al.: Heterogeneous neuronal firing patterns during interictal epileptiform discharges in the human cortex. Brain 133(6), 1668–1681 (2010)

[45] Truccolo, W., Donoghue, J.A., Hochberg, L.R., Eskandar, E.N., Madsen, J.R., Anderson, W.S., Brown, E.N., Halgren, E., Cash, S.S.: Single-neuron dynamics in human focal epilepsy. Nature neuroscience 14(5), 635–641 (2011)

[46] Khilkevich, A., Lohse, M., Low, R., Orsolic, I., Bozic, T., Windmill, P., Mrsic-Flogel, T.D.: Brain-wide dynamics linking sensation to action during decision-making. Nature 634(8035), 890–900 (2024)

[47] Rossant, C., Kadir, S.N., Goodman, D.F., Schulman, J., Hunter, M.L., Saleem, A.B., Grosmark, A., Belluscio, M., Denfield, G.H., Ecker, A.S., et al.: Spike sorting for large, dense electrode arrays. Nature neuroscience 19(4), 634–641 (2016)

[48] Watters, N., Buccino, A., Jazayeri, M.: Medicine: Motion correction for neural electrophysiology recordings. eneuro 12(3) (2025)

[49] Windolf, C., Yu, H., Paulk, A.C., Meszéna, D., Muñoz, W., Boussard, J., Hardstone, R., Caprara, I., Jamali, M., Kfir, Y., et al.: Dredge: robust motion correction for high-density extracellular recordings across species. Nature methods, 1–13 (2025)

[50] Hendrycks, D.: Gaussian error linear units (gelus). arXiv preprint 1606.08415 (2016)

[51] Belcak, P., Wattenhofer, R.: Fast feedforward networks. arXiv preprint 2308.14711 (2023)

[52] Hill, D.N., Mehta, S.B., Kleinfeld, D.: Quality metrics to accompany spike sorting of extracellular signals. Journal of Neuroscience 31(24), 8699–8705 (2011)

[53] Radmanesh, M., Rezaei, A.A., Jalili, M., Hashemi, A., Goudarzi, M.M.: Online spike sorting via deep contractive autoencoder. Neural Networks 155, 39–49 (2022)

[54] Reynolds, D.: Gaussian mixture models. In: Encyclopedia of Biometrics, pp. 827–832. Springer, ??? (2015)

[55] McInnes, L., Healy, J., Astels, S., et al.: hdbscan: Hierarchical density based clustering. J. Open Source Softw. 2(11), 205 (2017)

[56] Sarfraz, S., Sharma, V., Stiefelhagen, R.: Efficient parameter-free clustering using first neighbor relations. In: Proceedings of the IEEE/CVF Conference on Computer Vision and Pattern Recognition, pp. 8934–8943 (2019)

[57] Castellano-Escuder, P., Zachman, D.K., Han, K., Hirschey, M.D.: Gaudi: interpretable multiomics integration with umap embeddings and density-based clustering. Nature Communications 16(1), 5771 (2025)

